# ADHD symptoms and their neurodevelopmental correlates in children born very preterm

**DOI:** 10.1101/804799

**Authors:** Anita Montagna, Vjaceslavs Karolis, Dafnis Batalle, Serena Counsell, Mary Rutherford, Sophie Arulkumaran, Francesca Happe, David Edwards, Chiara Nosarti

**Affiliations:** Centre for the Developing Brain, School of Biomedical Engineering & Imaging Sciences, King’s College London, London, United Kingdom; Oxford Centre for Functional MRI of the Brain (FMRIB Centre), University of Oxford, Oxford, United Kingdom; MRC Social, Genetic & Developmental Psychiatry Centre, Institute of Psychiatry, Psychology & Neuroscience, King’s College London, London, United Kingdom; Department of Child and Adolescent Psychiatry, Institute of Psychiatry, Psychology and Neuroscience, King’s College London, London, United Kingdom

## Abstract

This study investigated the association between attention-deficit/hyperactivity disorder (ADHD) symptomatology in preschool-aged children who were born very preterm (<32 weeks) and cognitive outcomes, clinical risk and socio-demographic characteristics. 119 very preterm children who participated in the Evaluation of Preterm Imaging Study at term-equivalent age were assessed at a mean age of 4.5 years. Parents completed the ADHD Rating Scale IV, a norm-referenced checklist that evaluates ADHD symptomatology according to diagnostic criteria, and the Behavior Rating Inventory of Executive Function-Preschool version. Children completed the Wechsler Preschool and Primary Scales of Intelligence and the Forward Digit Span task. Longitudinal data including perinatal clinical, qualitative MRI classification, socio-demographic variables and neurodevelopmental disabilities were investigated in relation to ADHD symptomatology. All results were corrected for multiple comparisons using false discovery rate. Results showed that although the proportion of very preterm children with clinically significant ADHD did not differ from normative data after excluding those with neurodevelopmental disabilities, 32.7% met criteria for subthreshold ADHD inattentive type and 33.6% for combined type, which was higher than the expected 20% in normative samples. Higher ADHD symptom scores (all) were associated with greater executive dysfunction (inhibitory self-control, flexibility, and emergent metacognition, corrected p<0.001 for all tests). Higher inattentive ADHD symptom scores were associated with lower IQ (ρ=-0.241, p=0.036) and higher perinatal clinical risk (more days on mechanical ventilation (ρ=0.206, p=0.025) and more days on parenteral nutrition (ρ=0.223, p=0.015)). Higher hyperactive ADHD symptom scores instead were associated with lower socio-economic status (ρ=0.278, p=0.002). These results highlight the importance of monitoring and supporting the development of very preterm children throughout the school years, as subthreshold ADHD symptoms represent risk factors for psychosocial problems and for receiving a future clinical diagnosis of ADHD.

## Introduction

Attention-deficit/hyperactivity disorder (ADHD) is a neurodevelopmental condition which affects 5-7% of children, depending on the source of information used to reach a diagnosis (1, 2). Children who were born very preterm (< 32 weeks of gestation) have a 2- to 3-fold increased risk of being diagnosed with ADHD compared to their term born peers (4-fold risk in those born at < 26 weeks) (3–7).

According to the fifth edition of the Diagnostic and Statistical Manual of Mental Disorders (DSM-V), ADHD is characterized by two main symptom presentations, inattention and hyperactivity-impulsivity (8). Although symptoms often co-occur, their expression can be unbalanced, leading to a clinical subdivision of the disorder into inattentive, hyperactive-impulsive and combined types. In general population samples the most common form of ADHD presentation is the combined type, but among very preterm born children the inattentive subtype has higher prevalence (9, 10). There is further evidence that very preterm children display higher levels of subclinical inattentive symptomatology compared to controls, which falls below diagnostic thresholds (11, 12).

ADHD symptoms in very preterm children exhibit specific cognitive correlates that are not observed in term-born children with ADHD (3), and research has suggested that behavioural symptoms of inattention in very preterm children could be completely accounted for by slow responses and impairments in visuo-spatial working memory (13). Preterm children with ADHD do not show the typical pattern of higher prevalence in males compared to females (14, 15), and they tend not to have co-occurring conduct disorder (15, 16). Socio-demographic and environmental risks show a weaker association with ADHD in very preterm compared to term-born children, while, and perhaps not surprisingly, perinatal medical and neurological factors significantly contribute to their vulnerability to develop the disorder (17). These converging strands of evidence have led to the hypothesis of a possible ‘purer’ biological aetiology of ADHD following very preterm birth (3), supporting the idea of multiple pathways to ADHD (18).

The majority of studies to date in very preterm samples have focused on school age, when ADHD is typically diagnosed. However, the importance of detecting early risk factors associated with ADHD is well recognised (19). Attention problems identified during the pre-school years may mark the beginning of escalating academic problems, school drop-out and psychopathology later in life (20); indeed attention could be regarded as a building block for the development of many other cognitive functions and as a prerequisite for learning (21).

Here we investigate ADHD symptoms, their cognitive and clinical correlates in preschool-aged children who were born very preterm. We hypothesised that higher ADHD symptoms in very preterm children would be associated with lower executive functions and intelligence, even in the absence of a clinical diagnosis. We further hypothesised that inattentive ADHD symptoms would be associated with perinatal clinical adversity rather than environmental risk.

## Methods

### Sample

Study participants were 119 very preterm born children who were recruited at birth in 2010-2013 as part of the Evaluation of Preterm Imaging Study (e-Prime Eudra: CT 2009-011602-42) from hospitals within the North and Southwest London Perinatal Network (22). Infants were eligible if born before 33 weeks gestational age and their mother was aged over 16 years and not a hospital inpatient; they were excluded if they had major congenital malformation, prior Magnetic Resonance Imaging (MRI), metallic implants, parents unable to speak English, or were subject to child protection proceedings. The original sample recruited studied 511 of 1831 eligible infants and accurately reflected the population of preterm infants. Full data on the trial are available (22). The current study was a convenience sample created by families who accepted invitations to undergo further examination and testing at pre-school age (see Table 1 for a description of children’s perinatal, socio-demographic and cognitive characteristics). Current study participants did not differ from those in the larger cohort in terms of gestational age (F = 3.42, p > 0.05) and birth weight z score (F = 0.013, p > 0.05). Written informed consent was obtained from children’s carer(s) following procedures approved by the National Research Ethics Committee (14/LO/0677), in accordance with The Code of Ethics of the World Medical Association (Declaration of Helsinki).

**Table 1.** Study participants’ perinatal, socio-demographic and cognitive characteristics.

|  |  |
| --- | --- |
| **Gestational age in weeks, median (range)** | **30.00 (25.00 – 32.00)** |
| **Birth weight in grams, median (range)** | **1305 (600 – 2600)** |
| **Males, No (%)** | 54 (45.4) |
| **Days on mechanical ventilation, median (range)** | 0 (0 – 21) |
| **Days on parenteral nutrition, median (range)** | 5 (0 – 54) |
| **Intrauterine growth restriction, N (%)** | 17 (14.3) |
| **Index of multiple deprivation quintile, N (%)** |  |
| **1 (Least Deprived)** | 23 (19.3) |
| **2** | 24 (20.2) |
| **3** | 28 (23.5) |
| **4** | 24 (20.2) |
| **5 (Most Deprived)** | 20 (16.8) |
| **Corrected age at assessment in months, mean (range)** | 53.78 (50.06 – 59.19) |
| **Mother's age leaving full time education, at birth, N (%)\*** |  |
| **18 years or younger** | 33 (27.7) |
| **19 years or older** | 82 (68.9) |
| **Unknown** | 4 (3.4) |
| **Qualitative MR rating\*** |  |
| **No perinatal brain lesions N (%)** | 35 (29.4) |
| **Minor perinatal brain lesions N (%)** | 64 (53.8) |
| **Major perinatal brain lesions N (%)** | 17 (14.3) |
| **Full scale intelligence quotient, mean (range)** | 106 (50 -147) |

\*\*Three MRI datasets were corrupted and could not be evaluated

### Perinatal clinical and socio-demographic variables

Perinatal clinical and socio-demographic data were collected with permission from the Standardized Electronic Neonatal Database and included gestational age at birth, sex, days of mechanical ventilation, days of parenteral nutrition, intrauterine growth restriction (diagnosed by the obstetric team at the hospital where antenatal care was provided), mother’s age when leaving full time education and Index of Multiple Deprivation (IMD) score, determined by parents’ postcode at the time of infant’s birth (http://imd-by-postcode.opendatacommunities.org), which provide a proxy for socio-economic status.

Infants received MRI at term-equivalent age on a Philips 3 Tesla (Philips Medical Systems, Best, The Netherlands) system sited on the neonatal intensive care unit using an eight-channel phased array head coil. T2-weighted images were acquired (TR 8,670 ms; TE 160 ms; flip angle 90°; slice thickness 2 mm; in plane resolution 0.86 × 0.86 mm). An experienced neonatal radiologist qualitatively rated the MRI scan of each infant and assigned an overall global score that described the clinical severity of brain abnormalities. Scores ranged from 2-0; 2 = major lesion, defined as cystic periventricular leukomalacia, >10 punctate white matter lesions, grade 3 or 4 germinal matrix haemorrhage; 1 = minor lesion, defined as all any other lesions; 0 = no lesion.

### Behavioural outcomes

The ADHD Rating Scale-IV (ADHD-RS-IV) (23) was used to measure behaviours associated with different ADHD presentations according to DSM-V diagnostic criteria (8). This 18-item norm-referenced questionnaire provides scores from two separate subscales (Inattention and Hyperactivity/Impulsivity) and a combined score. These scales have been shown to have strong psychometric properties in preschool children (23). Raw scores were transformed into z scores using age- and sex-matched normative means and standard deviations. Scores were defined as clinically significant when equal or greater than the 90^th^ percentile of the sex-matched normative score distribution. The 80^th^ percentile was also used to include sub-threshold symptoms.

### Neuropsychological outcomes

General intellectual development was assessed with the Wechsler Preschool and Primary Scales of Intelligence battery (WPPSI-IV) (24). Age corrected for prematurity was used to calculate scaled scores (25, 26). Phonological short term memory was assessed using the digit span forward test, using an adapted version of the Wechsler Intelligence Scale for Children IV digit span task (27). Inhibitory Self-Control Index (Inhibition and Emotional Control scores), Flexibility Index (Shifting and Emotional Control scores) and Emergent Metacognition Index (Working Memory and Planning/organisation scores) were obtained from the Behavior Rating Inventory of Executive Function-Preschool version (BRIEF-P) (28). Scores are reported as T-scores, with higher scores indicating greater executive dysfunction.

### Neurodevelopmental Disabilities

Motor impairment was determined on the basis of scores below 2 standard deviations on the Gross Motor Scale of the Bayley Scales of Infant Development-III (29) obtained at 18-22 months of age. Neuromotor development was further evaluated at current assessment by a trained researcher and reviewed by an experienced paediatrician. The assessment consisted of a brief neurological examination to confirm the presence or absence of cerebral palsy. Sensory deprivation, speech problems and psychiatric diagnoses (or suspected clinical significance together with the child waiting for a specialist’s opinion) were not directly assessed and were based on parental reports. Psychiatric diagnoses included autism spectrum disorder (ASD) and global developmental delay (GDD). In case of reported problems, the child’s General Practitioner was contacted, and medical notes were accessed after obtaining caregivers’ consent.

Children with at least one of the following were regarded as having a neurodevelopmental disability (NDD): neuromotor impairment (assessed at 18-22 months and confirmed by subsequent diagnosis of cerebral palsy), diagnosis of ASD and/or GDD, sensory or/and speech problems.

### Data analysis

SPSS version 22.0 (SPSS Inc, Chicago, IL) and Matlab (MathWorks Inc, Natick, MA, U.S.A) were used for analyses. Binomial tests were used to compare the proportion of very preterm children with clinically significant (and subclinical) levels of behavioural problems against age-matched UK population norms. Analyses were run before and after exclusion of children with NDD.

To assess the prevalence of inattention and hyperactivity/impulsivity, a data-driven clustering procedure was also used on the ADHD-RS-IV scores to fractionate the sample into groups, defined by the imbalance between their inattentive and hyperactive/impulsive symptoms, similar to (30). An Expectation-Maximisation (EM) algorithm fitting a Gaussian mixture model was used to identify relatively homogeneous groups of cases. Coordinates for centroids were initialized by running k-means clustering, for which centroid coordinates were initialised at random.

The associations between ADHD symptoms, IQ and neurocognitive skills (phonological short-term memory, inhibitory self-control, flexibility and emergent metacognition) were investigated using non-parametric Spearman’s rank-order correlation. Analyses were run after exclusion of children with NDD and p values were always corrected for multiple comparisons using a false discovery rate (FDR) approach controlling alpha error to 5% (31). Spearman’s correlations were used to investigate the relationship between ADHD symptoms and perinatal clinical variables and socio-economic status. Sex differences in ADHD scores were investigated using the Mann-Whitney U test. After testing for the presence of significant outliers (calculated using an interquartile range), variance homogeneity among groups (Levene’s test of Equality of Error Variances), and approximately normal distribution in the different groups, Kruskal-Wallis tests were used to test the association between severity of neonatal brain abnormalities and ADHD symptoms.

Comorbidities between NDD and behavioural problems were defined by having NDD plus ADHD symptoms above the 90^th^ centile of the sex-matched normative score distribution.

## Results

### ADHD Symptomatology

The proportion of very preterm children with clinically significant inattentive (20.2%), hyperactive/impulsive (16.0%) and combined symptoms (16.8%) was higher than the expected 10% in normative samples of predominantly term-born individuals (after FDR correction; see Table 2). However, after exclusion of children with NDD, who represented 10.1% of the sample, the proportion of very preterm children with clinically significant ADHD symptoms did not differ from the population norm. In fact, comorbidity between NDD and clinically significant ADHD symptoms was high: 66.7% of children with NDD had inattention symptoms above the 90^th^ centile cut-off, 25.0% had hyperactivity/impulsivity and 58.3% had combined symptoms. When using the 80^rd^ percentile cut-off and excluding children with NDD, 32.7% of very preterm children had inattentive symptoms and 33.6% had combined symptoms, which was higher than the expected 20% in normative samples.

**Table 2.**
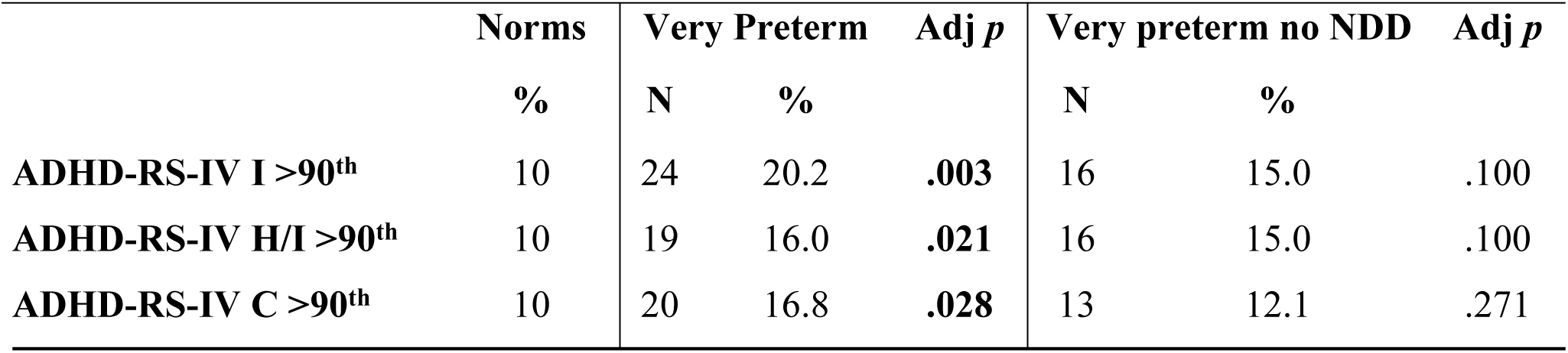
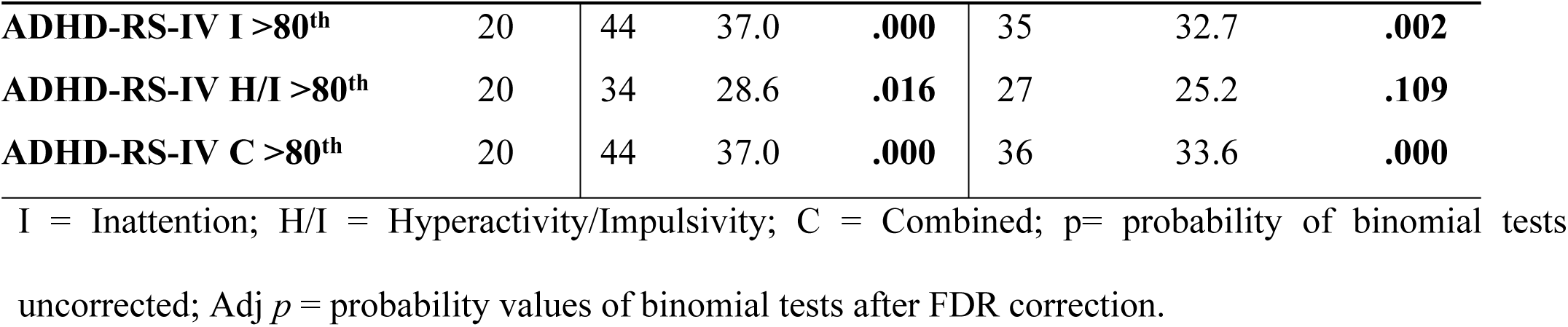
Rates of clinical (>90th centile cut-off) and subthreshold (>80th centile cut-off) ADHD symptoms in very preterm children before and after exclusion of those with NDD.

K-means clustering identified four ADHD groups: 13 children with predominantly inattentive symptoms (10.9%), 12 with high levels of both inattentive and hyperactive/impulsive symptoms (10.1%), 44 with medium levels of both symptoms (37%), 50 with low levels of both symptoms (42%) and no child with mainly hyperactive/impulsive symptoms.

**Figure 1.**
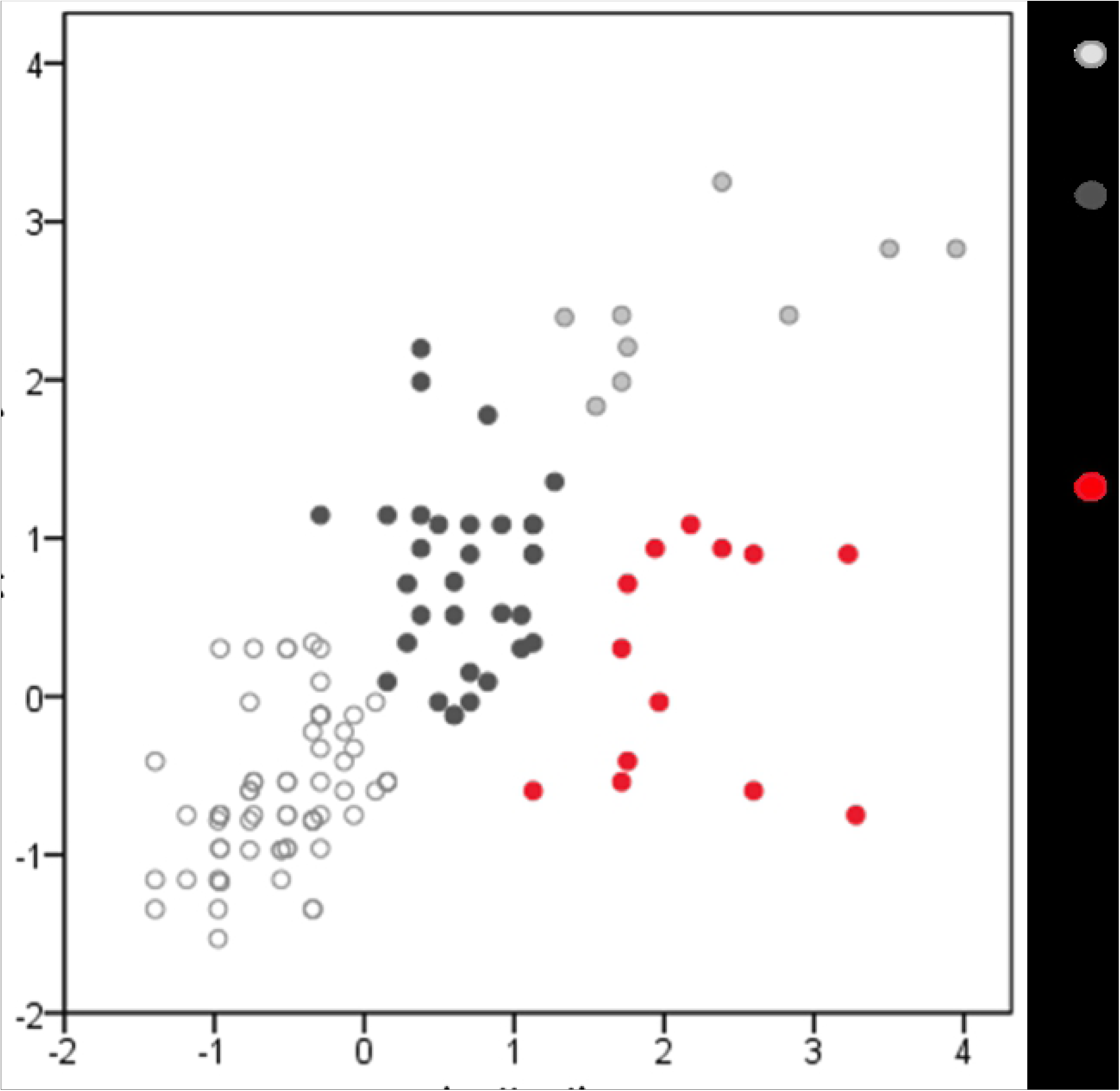
Visual representation of inattentive and hyperactive symptoms in all study participants after exclusion of children with NDD.

### Sex differences in ADHD symptomatology

Mann-Whitney tests showed no sex differences in inattentive (U = 1.429; *p* = 0.082), hyperactive/impulsive (U = 1.656; *p* = 0.598) and combined ADHD symptom scores (U = 1.536; *p* = 0.243), even after exclusion of children with NDD.

### Associations between ADHD symptomatology and cognitive variables

After excluding children with NDD, higher inattentive ADHD symptom scores were associated with lower full-scale IQ (ρ = −0.245, *p* = 0.011) but not with hyperactive and combined ADHD symptom scores (ρ = −0.135, *p* = 0.168 and ρ = −0.190, *p* = 0.050, respectively). Higher ADHD symptom scores (all) were associated with poorer inhibitory self-control, flexibility and emergent metacognition scores, as shown in Table 3.

**Table 3.** Associations between ADHD-RS-IV symptom subtype scores and cognitive outcomes after exclusion of children with NDD (n=107).

|  | Digit Span |  | Inhibitory<br>Self-Control |  | Flexibility |  | Emergent<br>Metacognition |  |
| --- | --- | --- | --- | --- | --- | --- | --- | --- |
| | $\rho$ | $p$ | $\rho$ | $p$ | $\rho$ | $p$ | $P$ | $p$ |
| **Inattentive** | -0.064 | 0.516 | 0.542 | <0.001* | 0.481 | <0.001* | 0.735 | <0.001* |
| **Hyperactive** | -0.140 | 0.150 | 0.554 | <0.001* | 0.459 | <0.001* | 0.582 | <0.001* |
| **Combined** | -0.101 | 0.299 | 0.586 | <0.001* | 0.505 | <0.001* | 0.718 | <0.001* |

Spearman correlation; $\rho$ =Rho; $p$ = probability values; \* significant after FDR correction.

### Associations between ADHD symptomatology, perinatal clinical risk and socio-demographic variables

Higher inattentive ADHD symptom scores were associated with more days on mechanical ventilation (ρ(119) = 0.196, *p* = 0.032) and more days on parenteral nutrition (ρ(119) = 0.222, *p* = 0.015). All results survived FDR correction. There was no evidence of differences in ADHD symptom scores according to severity of perinatal brain lesions (inattentive, *H* = 3.038, p = 0.219; hyperactive, *H* = 1.646, p = 0.439; combined, *H* = 1.858; p = 0.395).

Hyperactive/impulsive and combined ADHD symptom scores were correlated with higher IMD scores which reflect lower socio-economic status (ρ(119) = 0.259, *p* = 0.004 and ρ(119) = 0.198, *p* = 0.031, respectively). Children belonging to the 5^th^ IMD quintile, reflecting the most deprived socio-economic group, had a z score of 0.438, while those children belonging to the 1^st^ IMD quintile, reflecting the least deprived socio-economic group, had a z score of −0.433. After FDR correction, these associations remained unaltered. Similar findings persisted when children with NDD were excluded from each analysis.

## Discussion

Although the proportion of very preterm children with clinically significant ADHD did not differ from normative data after excluding those with neurodevelopmental disabilities, 32.7% met criteria for subthreshold ADHD inattentive type and 33.6% for combined type, which was higher than the expected 20% in normative samples. These results highlight the importance of monitoring and supporting the development of very preterm children throughout the school years, as subthreshold ADHD symptoms have been associated with an increasing risk of psychosocial problems and of receiving a future diagnosis of the clinical disorder (32–34).

In addition to the high subthreshold ADHD inattentive symptom prevalence, our results showed that inattention was a core deficit in this sample of very preterm children. K-means clustering identified a group of very preterm children with predominantly inattentive or combined symptoms, but did not identify a subgroup of children with predominantly hyperactive/impulsive ADHD symptoms. This is in contrast with what described by Sanefuji and colleagues (2016) who similarly used k-means clustering in a sample of 165 children diagnosed with ADHD and were able to parse children based on the three typical ADHD clinical presentations (30).

Psychiatric risk in extremely preterm children has been found to be higher in those with coexisting neurodevelopmental disabilities (35), although neurodevelopmental disabilities have been also associated with psychopathology irrespective of very preterm birth (36). In this study, up to two thirds of children with clinically significant ADHD-inattentive symptoms had comorbid neurodevelopmental disabilities, as did one quarter of those with clinically significant ADHD-hyperactive/impulsive symptoms. In atypically developing children a complex relationship exists between cognitive outcome, biological and environmental factors (37), and here we attempted to elucidate the association between ADHD symptomatology, cognitive and environmental outcomes, cerebral injury and perinatal clinical risk.

Inattentive ADHD symptoms were associated with lower IQ, which has been studied as a longitudinal predictor of educational outcomes (20). There is a strong phenotypic association between ADHD and lower IQ which has been attributed to shared genes (38) and previous research suggested that this association may be particularly strong for inattentiveness (39). It is therefore possible that preterm birth may share common genetic influences with IQ and academic achievement (40).

Previous findings in older preterm children suggested that a core executive difficulty underlined their inattentive problems (13, 41, 42), while the current study reported an association between higher ADHD symptoms (all types) and executive scores. These included inhibitory self-control, flexibility and emergent metacognition, which refers to the process of active control over one’s own cognition, encompassing processes involved in self-appraisal and self-management (43) and have been studied as core features of ADHD (44, 45). The lack of ADHD symptom specificity in relation to executive functions observed here could be explained by methodological issues associated with modelling executive function early in development (46). More studies are therefore required to further investigate these associations.

The current finding of a positive correlation between inattentive ADHD symptoms and perinatal medical complications suggests that preterm-born pre-schoolers display a phenotypic profile that differs from that observed in the general population, with aetiological underpinnings in the clinical risk associated with preterm birth. This was assessed using markers of general infant health: number of days of parenteral nutrition (reflecting gut failure) and number of days spent on invasive ventilation via an endotracheal tube (reflecting respiratory failure). In a larger sample of infants the current study participants were drawn from, these two risk factors were independently associated with lower fractional anisotropy values throughout the white matter at term equivalent age, reflecting alterations in brain development (47). These results suggest that although in this study ADHD symptomatology did not differ according to qualitatively-rated severity of perinatal brain lesions (unlike previous findings in 11 years old children who were born extremely preterm (5)), future studies should include measures of brain structure and function to further characterise the association between clinical risk and brain development in order to better understand possible causative pathways leading to inattention in preterm children.

The idea that perinatal events can be implicated in the aetiology of psychiatric disorders has long since been proposed (48) and several studies have supported this hypothesis by showing that preterm birth may act as independent predictor of later psychopathology (49). The causal pathway to later psychiatric disorder could be interpreted in the context of the neurodevelopmental sequelae of preterm birth (50), although little is known about the neurobiological mechanisms leading to different ADHD subtypes. In children with ADHD a double dissociation has been shown between resting state functional connectivity in specific networks associated with inattentiveness and hyperactivity-impulsivity, indicating that different brain alterations may characterise ADHD subtypes (30).

The early identification of a preterm profile characterised by inattentive ADHD symptomatology associated with lower IQ and perinatal clinical risk suggests the need for early screening and educational interventions aimed at improving specific attention problems in very preterm children, many of whom exhibit academic difficulties at school age and are three times more likely to have special educational needs than their term-born peers (51). Attention is a crucial requirement in school settings as it allows the child to engage with classroom education activities. A strong relation between attention problems and academic achievement has been shown in previous studies (even after accounting for IQ) (20) and attention is considered one of the major areas of competence, which contribute to determine a successful transition to school (“school readiness”) (52). The use of easy and quick screening questionnaires, such as the ADHD-RS-IV, could help to identify potential targets for the prevention of the escalation of academic problems that have been associated with inattentiveness (53). This is especially important, given that attention problems tend to be more persistent in time and more difficult to be detected in the school setting compared to hyperactive/impulsive behaviours (10, 20). Moreover, preschool-age inattention is associated with other areas of difficulty, including communication and socio-emotional skills (52). Therefore, the allocation of additional support for very preterm children before school entry could enable them to enter school on a more equal footing with their peers.

Environmental variables including socio-economic status have been studied as risk factors for ADHD (54), although the majority of published studies have not differentiated between ADHD symptoms subtypes. Our results showed that children from socially disadvantaged families had increased hyperactive/impulsive but not inattentive ADHD symptoms. These findings and are in line with others, showing selective associations between hyperactivity/impulsivity and environmental variables, i.e. an unsupportive home environment (55). The relationship between socioeconomic disadvantage and ADHD is likely to be intricate and may be mediated by other factors that are associated with lower socio-economic status, such as lower parental education and a less stimulating home environment (56).

Limitation of this study include the lack of a locally matched control group. This could have influenced the results, possibly underestimating the level of problems experienced by very preterm children (57). Strengths of this study include the use of a wide battery of tests and of a questionnaire aimed at exploring ADHD subtypes, the use of a strict statistical procedure to correct for multiple testing, the inclusion of children with NDD in the analysis, the use of data driven together with theory driven approaches, the use of longitudinal information together with cross-sectional data.

In summary, this study shows that already during preschool age very preterm children are at increased risk of sub-threshold ADHD symptoms, especially the inattentive type. Inattentive ADHD symptoms were associated with intellectual functioning at time of assessment, and with perinatal clinical risk, supporting the hypothesis of a possible neurobiological origin of ADHD. In light of these findings, further investigation of early inattentive symptoms may support the development of effective interventions and provide new models for understanding the neurofunctional trajectories leading to ADHD in children born very preterm.

## List of Abbreviations and Acronyms

ADHD: Attention-deficit/hyperactivity disorder
MRI: magnetic resonance imaging
NDD: Neurodevelopmental disability

## Acknowledgements

We are grateful to the families who generously took part in this research. This work was supported by the Medical Research Council (UK), the National Institute for Health Research (NIHR) comprehensive Biomedical Research Centre award to Guy’s & St Thomas’ National Health Service (NHS) Foundation Trust in partnership with King’s College London and King’s College Hospital NHS Foundation Trust. We thank all e-Prime investigators for their contribution to the study. This study used perinatal data acquired during independent research funded by the NIHR under its Programme Grants for Applied Research Programme (Grant Reference No. RP-PG-0707-10154). The views expressed are those of the authors and not necessarily those of the NHS, the NIHR, or the Department of Health.

